## Supplementary files for "Multi-scale model suggests the trade-off between protein and ATP demand as a driver of metabolic changes during yeast replicative ageing"

#### Supplementary Figures

Barbara Schnitzer *[1,2], Linnea Österberg *[3], Iro Skopa [1,2], Marija Cvijovic [1,2]

* Authors contributed equally
[1] Department of Mathematical Sciences, Chalmers University of Technology, Gothenburg, Sweden
[2] Department of Mathematical Sciences, University of Gothenburg, Gothenburg, Sweden
[3] Department of Biology and Biological Engineering, Chalmers University of Technology, Gothenburg, Sweden

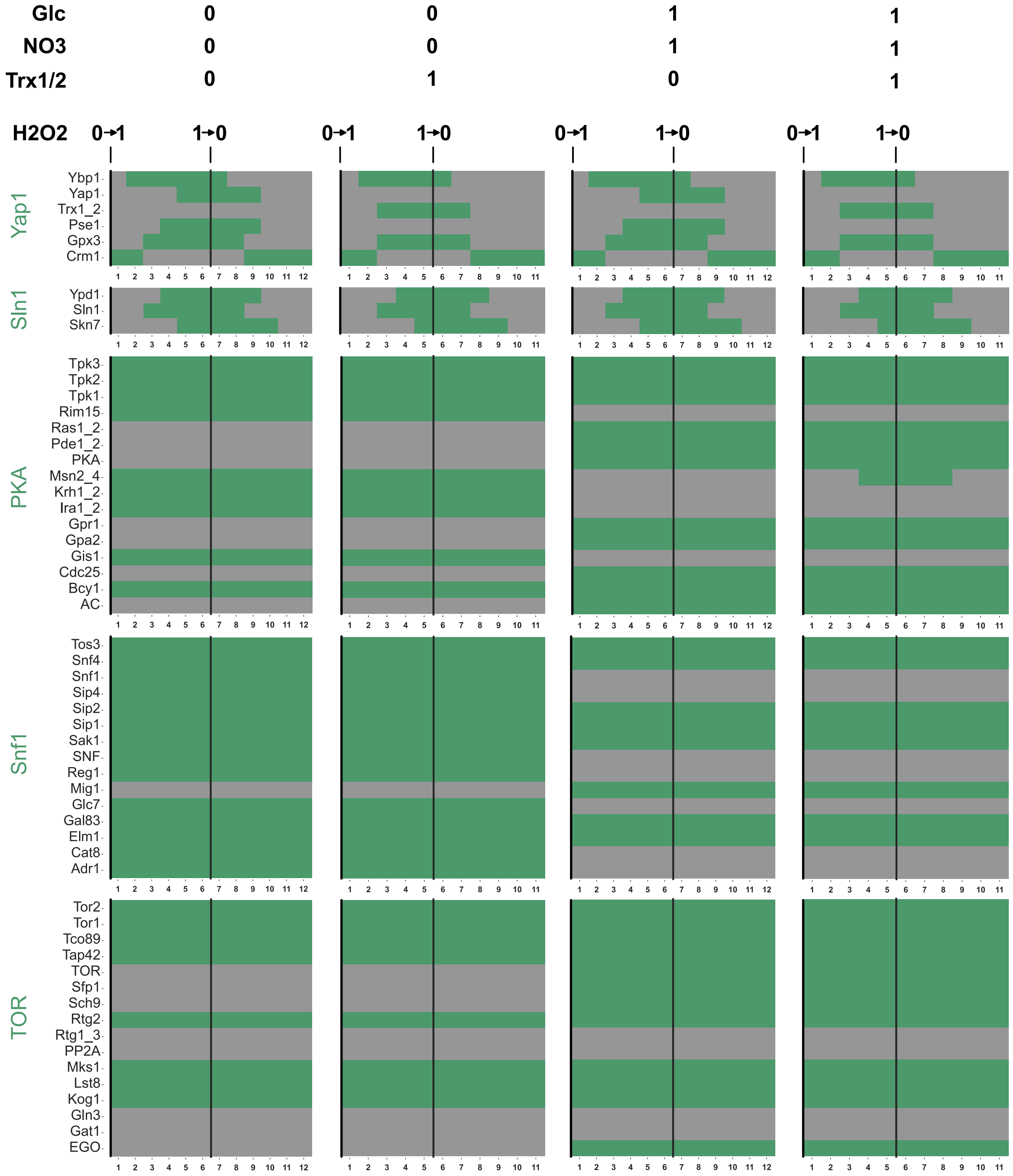

**Figure S1: Validation of Boolean model extension** Boolean model states when turning on $(0 \rightarrow 1)$ and off $(1 \rightarrow 0)$ hydrogen peroxide ($H_2O_2$) depending on glucose and nitrogen availability and the presence of thioredoxins (Trx1/2). $H_2O_2$ eventually triggers active transcription factors Skn7 and Yap1, the latter only if Trx1/2 is not active. Active Trx1/2 inhibits Yap1 activation and activates Msn2/4.

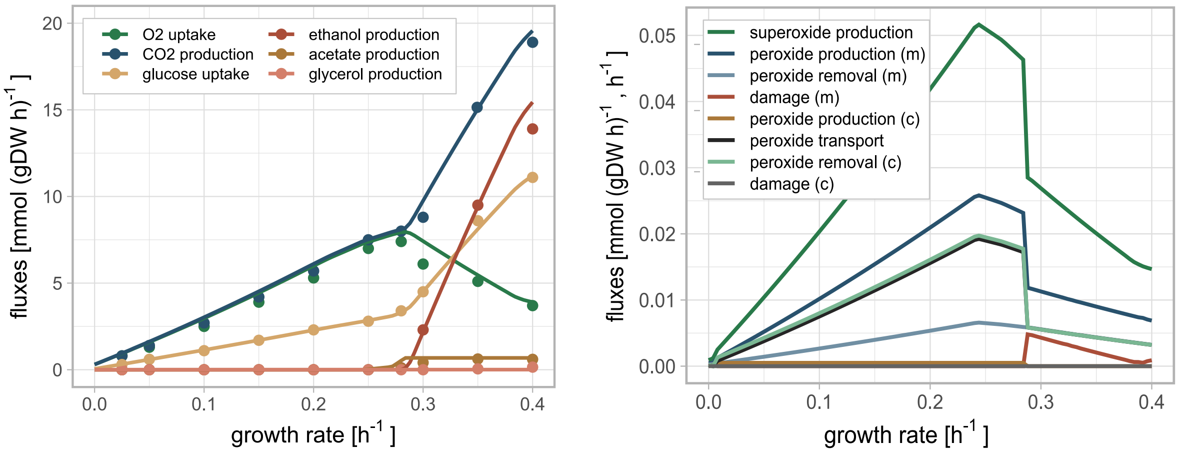

**Figure S2: Validation of the ecFBA model extension** Simulation of the chemostat experiment [46] using the extended regulated enzyme-constrained metabolic model. The left panel shows exchange fluxes in the model (solid lines) compared to the experimental data (dots). The right panel shows damage-related fluxes in the model, where chemostat data is not available.

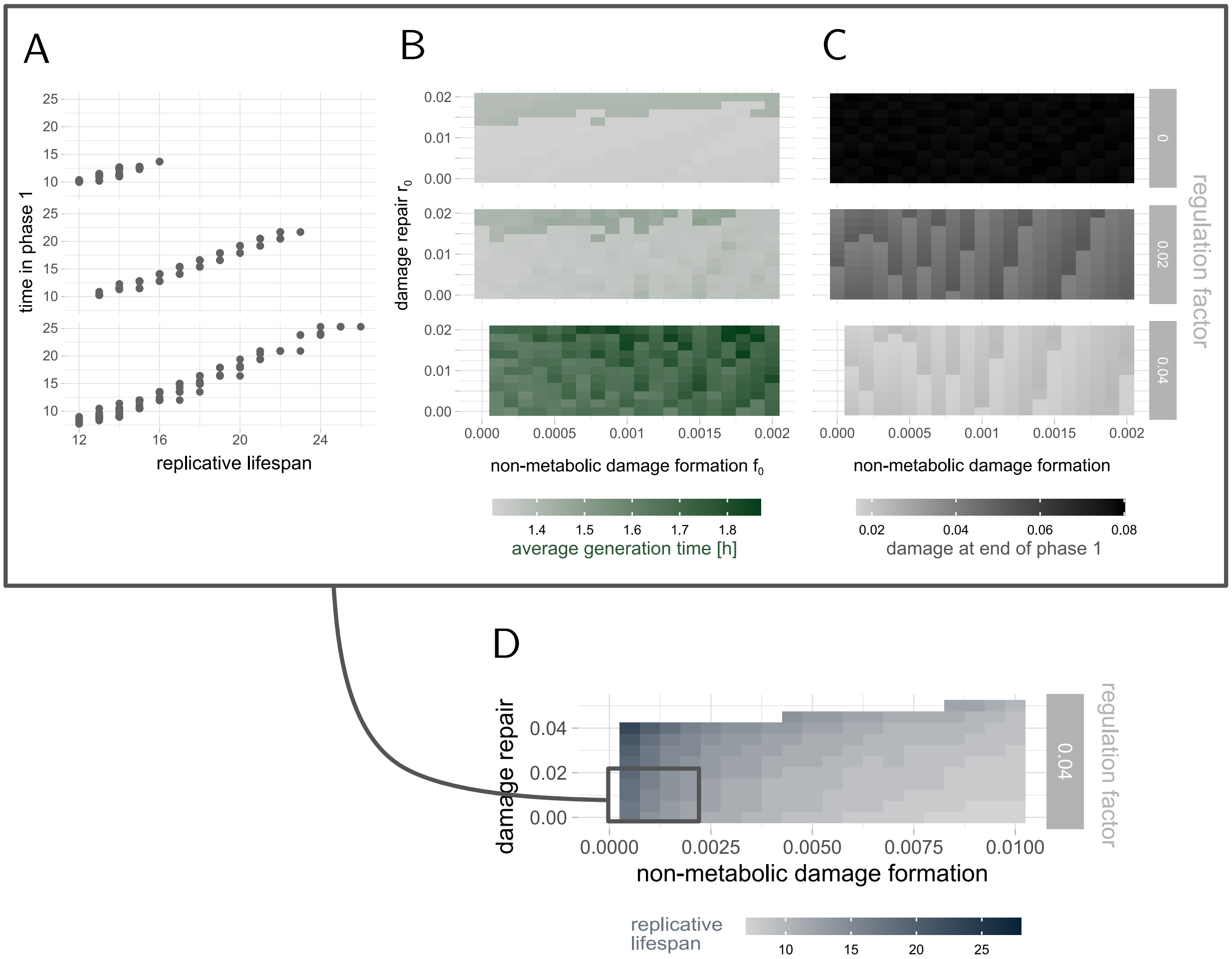

**Figure S3: Complementary simulations of yeast cells** for varying damage repair $r_0$ and non-metabolic damage formation rates $f_0$. (A) Correlation between the time cells spend in phase I and their replicative lifespan. (B) Average generation times. (C) Fraction of damaged protein content at the end of phase I. (D) Replicative lifespans. (A)-(C) are coming from the same data set as Fig 2C. The grid in (D) is further zoomed out. If there is no colour it means that the cell did not stop dividing in the simulation time, since repair is so efficient that it overcomes damage formation and retention.

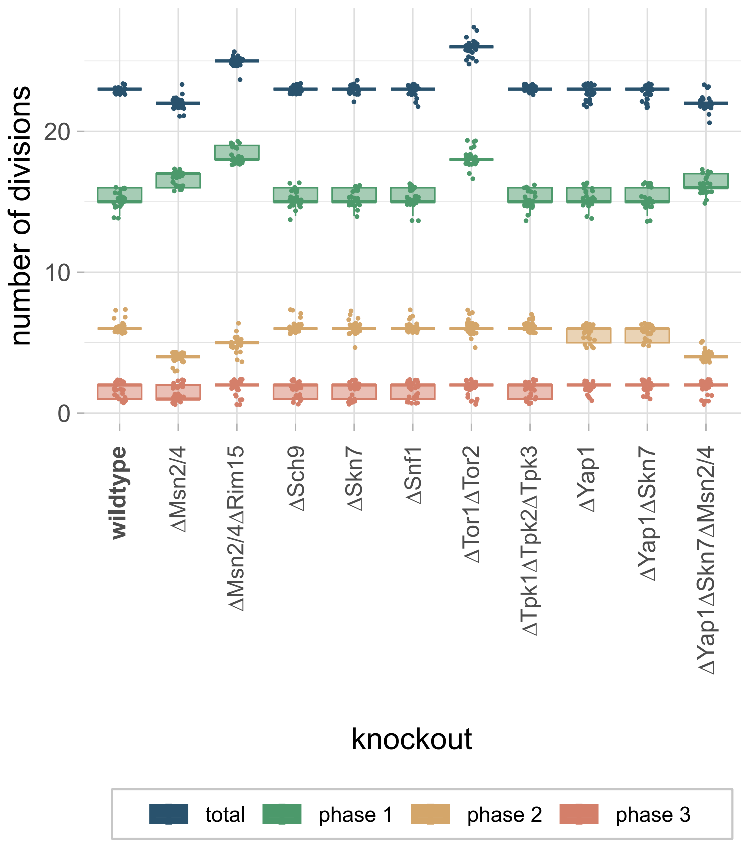

**Figure S4: Effect of signalling protein knockouts on phases** Split of the number of divisions over the metabolic phases for knockouts of signalling proteins in the different pathways of the Boolean model for the cells in Fig 3B. The distributions are based on 29 wildtype parameter sets with $f_0 \leq 5 \cdot 10^{-4}$ and $r_0 \leq 2 \cdot 10^{-2}$ that lead to 23 divisions (from data in Fig 2A). A lifespan increase is mostly achieved by prolonging phase I.

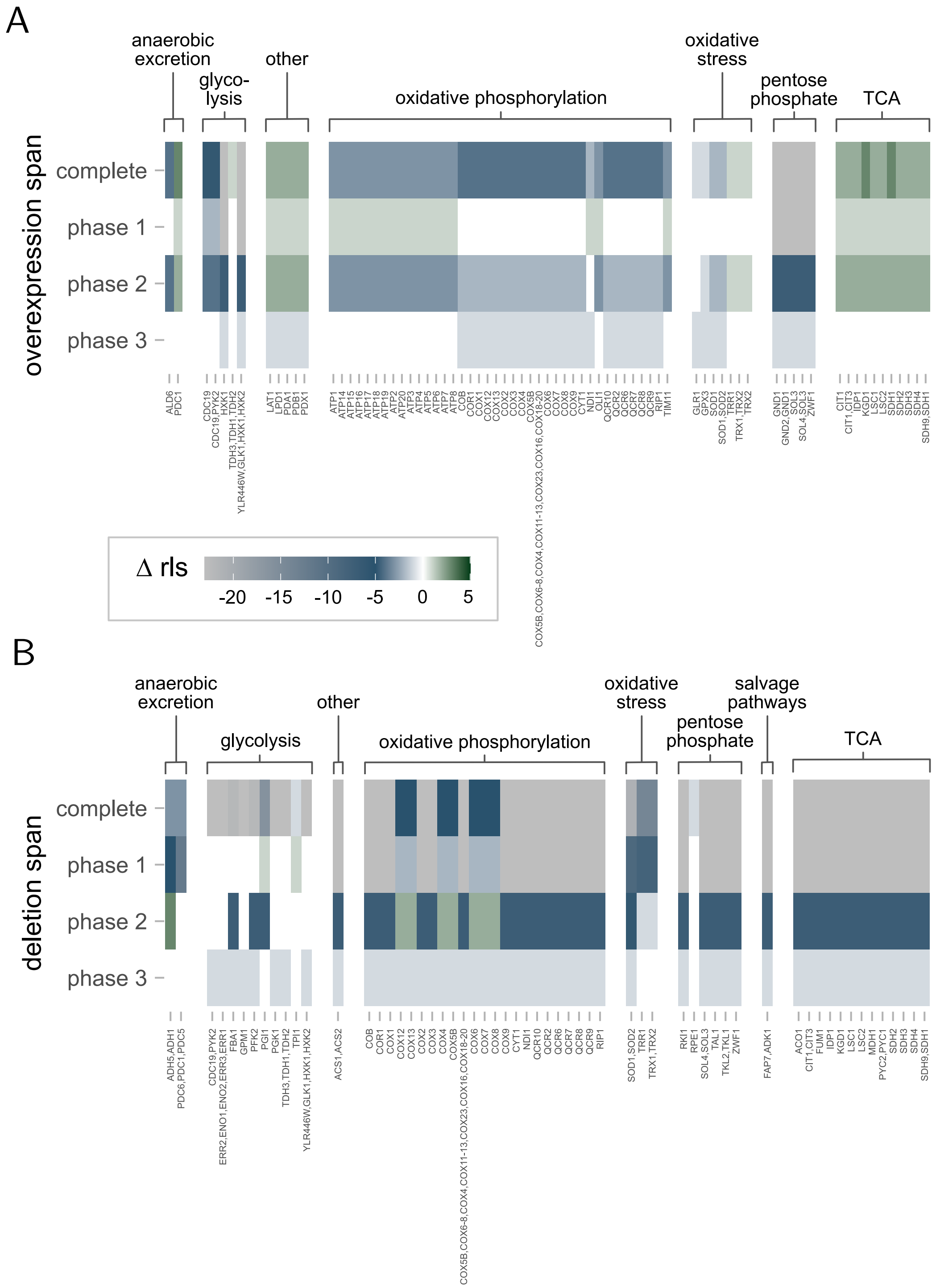

**Figure S5: Effect of enzymes perturbations on phases** Absolute changes in the number of divisions in relation to a wildtype cell (23 divisions) for overexpressions (A) or deletions (B) of single enzymes or isoenzyme combinations (140 + 23 cases) in different phases in the metabolic model. The simulations are based on $f_0 = 0.0001$ and $r_0 = 0.0005$ (as in Fig 2B and Fig 4). Enzymes that do not lead to any differences are excluded in this plot. Perturbation in a specific phase can have a different effect on the lifespan than the same perturbation over the whole life.

### Multi-scale model suggests the trade-off between protein and ATP demand as a driver of metabolic changes during yeast replicative ageing
### Supplementary text 1: Model details

Barbara Schnitzer [*1,2], Linnea Österberg [*3], Iro Skopa [1,2], Marija Cvijovic [1,2]

March 7, 2022

[*] Authors contributed equally
[1] Department of Mathematical Sciences, Chalmers University of Technology, Gothenburg, Sweden
[2] Department of Mathematical Sciences, University of Gothenburg, Gothenburg, Sweden
[3] Department of Biology and Biological Engineering, Chalmers University of Technology, Gothenburg, Sweden

**3 Dynamical model of growth, cell division and damage accumulation** **4**

### 1 Boolean model of cellular signalling

#### 1.1 Theoretical background

Vector-based Boolean models are powerful tool to understand the topology of a network. They are generally based on logical arguments and moreover parameter-free. Given a network, each of its $N$ components is represented by a p-dimensional vector of binary states $\mathbf{p}_i \in \{0,1\}^k$, $i = 1..N$, that represent chosen properties. The state of each component $i$ can be altered based on all other states by Boolean rules or functions $\mathcal{B}$ : $(\mathbf{p}_1, \mathbf{p}_2, .., \mathbf{p}_N) \to \mathbf{p}_i$. In each iteration all Boolean functions $\mathcal{B}_j$, $j = 1..M$, are applied synchronously to the states to generate updated states for the next step. Eventually, the system can end up in a logical steady state where the states of all components remain constant.

This formalism can be used to understand signalling events in cells in order to find the eventual activation of transcription factors. Here, we used a vector-based Boolean model of the nutrient signalling pathways Snf1, Tor and PKA that was previously published [1] and extended it further with the oxidative stress signalling pathways Yap1 and Sln1, as well as crosstalk to Msn2/4 that is also part of the nutrient signalling pathway PKA (Fig 1A). Each component in the network is represented by the properties presence, phosphorylation, oxidation and specific activity ($k = 4$). The Boolean rules are based on an extensive literature review to describe the signalling events, and correspond to simple if-statements, such as: IF protein X is present and phosphorylated AND protein Y is present and oxidised then protein Z gets phosphorylated. The activity property is especially important in our setting and is interpreted as being in the nucleus and interacting with the DNA, which can lead to expression or repression of genes. In particular, one can therefore investigate how perturbed input signals regarding nutrient availability and stress can influence the resulting transcription factor activity. Moreover, Boolean models generally allow to find logical gaps in the network [2, 3].

#### 1.2 Addition of oxidative stress signalling

In total, we added 9 new components and 13 new rules, including 1 crosstalk reaction, to the existing model of nutrient signalling [1] to account for oxidative stress signalling by the Yap1 and the Sln1 pathway.

**Yap1**
Yap1 mediates ROS stress signalling by sensing of $H_2O_2$ mediated trough the peroxidin Gpx3[4, 5]. In the presence of $H_2O_2$ Gpx3 together with Ybp1 facilitates the formation of active Yap1 [4, 5] that will accumulate in the nucleus where it induces it's gene targets including SOD1, GSH1, GPX2, TRX2 and TSA1 [6, 7]. The pathway returns to it's reduced state when reduced by thioredoxin, which is also a target gene of Yap1 [8].

**Sln1**
The Sln1 pathway is associated with osmoregulatory response, where Sln1 regulates Ypd1 phosphorylation. Ypd1 akts on the Ssk1 and on the transcription factor Skn7 [9, 10]. The role of this pathway in ROS regulation is not elucidated, but there are increasing reports of it's association to oxidative stress response where Skn7 plays a role, alone and in connection with Yap1 [9, 10].

**Crosstalk to nutrient signalling**
Msn2 and Msn4 are central stress regulators, targeting an number of oxidative response genes. The activation in respons to oxidative stress is mediated trough the thioredoxins Trx1 and Trx2[11]. Knockouts of Msn2 or Msn4 exhibit hypersensitivity to $H_2O_2$ and the response is only partially overlapping with that of Yap1 and Skn7 [12].

### 2 Enzyme-constrained flux balance analysis of the metabolic network

#### 2.1 Theoretical background

Generally, in flux balance analysis (FBA) [13–15] chemical reactions in the network are represented by a stoichiometric matrix $S$. Assuming that each component can only be used as much as it is produced, the system is naturally constraint by this mass balance requirement. Mathematically, the fluxes $\mathbf{v}$ through the network have to satisfy $S\mathbf{v} = 0$. Given an objective function, that can be an individual flux or a combination of several fluxes, all fluxes can be optimised accordingly. Biologically, examples for objective functions are maximal growth, minimal nutrient uptake or maximal growth yield. The optimal solution can be found by solving the linear program in (1).

$$\begin{aligned} \text{optimise} \quad & z = \mathbf{c}^T \mathbf{v} \\ s.t. \quad & S\,\mathbf{v} = \mathbf{0} \\ & \mathbf{v}_{min} \leq \mathbf{v} \leq \mathbf{v}_{max}, \end{aligned} \tag{1}$$

with $\mathbf{c}$ defining the coefficients of the fluxes in the objective function. Furthermore, $\mathbf{v}_{min}$ and $\mathbf{v}_{max}$ are general lower and upper bounds on the fluxes.

Enzyme-constrained FBA (ecFBA) [16, 17] is an extension of the traditional FBA, incorporating enzymes as components that are required for catalysing certain reactions. Each enzyme $e_i$ that is used is drawn from an enzyme pool $e_{pool}$ and is consumed in one or more reactions with a stoichiometric coefficient inversely proportional to its respective turnover number $k_{cat}$. The enzyme pool is itself restricted by the total amount of proteins $P_{tot}$ in the cell. The new additional constraints in the optimisation problem are stated in (2).

$$\begin{aligned} s.t. \quad & -\sum_j \frac{n^{ij}}{k_{cat}^{ij}} v_j + e_i = 0, \qquad \forall i \\ & -\sum_i MW_i e_i + e_{pool} = 0 \\ & \mathbf{e}_{min} \leq \mathbf{e} \leq \mathbf{e}_{max} \\ & 0 \leq e_{pool} \leq \sigma f P_{tot}, \end{aligned} \tag{2}$$

with $n^{ij}$ being the number of enzymes $i$ that are needed to catalyse reaction $j$. In most cases $n^{ij}$ equals 0 or 1, but can in exceptional cases of enzyme complexes be higher. Further, $MW_i$ are the molecular weights of the enzymes, $f$ corresponds to the fraction of the total protein mass covered by the enzymes in the model and $\sigma$ to the saturation factor of the enzymes. Similar to before, $\mathbf{e}_{min}$ and $\mathbf{e}_{max}$ are general lower and upper bounds on the enzyme usages. Typically, each optimisation is followed up by a second optimisation that picks the solution with a minimal sum of all fluxes and enzyme usages (parsimonious FBA). EcFBA has been shown to improve the predictive power in comparison to the traditional FBA [1, 17].

Note that fluxes typically have the unit $[mmol(gDWh)^{-1}]$ or $[h^{-1}]$, while enzyme usages are measured in $[mmol(gDW)^{-1}]$ and protein content in $[g(gDW)^{-1}]$.

#### 2.2 Addition of damage producing reactions

In this work, we make use of a previously published ecFBA model of the central carbon metabolism [1, 16] and incorporated new chemical reactions that produce reactive oxygen (ROS) and nitrogen species (RNS) (Fig 1B).

The new reactions are based on the fact that while cells produce energy in the mitochondria about 0.2-2% electron leak from the electron transport chain (ETC) [18]. Complex 3 in the ETC can be responsible for some of those electrons, while most of them escape from complex 1 [18–20]. The major downstream ROS and RNS reactions that are caused by the free electrons are summarised in the following according to [21–27]. When electrons react with oxygen ($O_2$) the negatively charged superoxide ($O_2^-$) is produced. $O_2^-$ can be transformed to $H_2O_2$ via superoxide oxidoreductase (SOD1, SOD2). $H_2O_2$ can be transformed back to water by glutathione peroxidase (GPX1-3). In that reaction glutathione disulfide gets glutathione. To transform back glutathione to glutathione disulfide the enzyme glutathione oxidoreductase (GLR1) is needed. Similar reactions happen for thioredoxin instead of glutathione, using thioredoxin peroxidase (TRX1-3) and reductase (TRR1-2). In addition, $H_2O_2$ transforms to $OH^{\cdot}$ via Fenton- and Haber-Weiss reactions with iron cations as mediators. $OH^{\cdot}$ can also be indirectly produced by $OONO^-$, encompassing several reactions that besides

others convert the nitric oxide radical $NO^{\cdot}$ to $NO_2^{\cdot}$. In this simplified pathway the major cause of damage is the hydroxyl radical ($OH^{\cdot}$) that can oxidise proteins and make them dysfunctional.
In addition, we introduced a non-growth associated ATP cost (NGAM) to the model, $NGAM(t) = \frac{D(t)}{P(t)+D(t)} \cdot NGAM_{max}$., with the $NGAM_{max}$ as in [28].
In total, it resulted in 52 new reactions and 41 new components including 13 new enzymes in the ecFBA model compared to [1]. Necessary $k_{cat}$ values were adopted from the consensus yeast metabolic model [28].

### 3 Dynamical model of growth, cell division and damage accumulation

To describe the protein damage accumulation over time, we make use of an ordinary differential equation (ODE) model, that is based on three forces: damage formation, damage repair and cell growth. The biomass of a cell $M$ $[gDW]$ follows a simple linear ODE with a time-dependent growth factor $g(t)$.

$$\frac{\mathrm{d}M(t)}{\mathrm{d}t} = g(t)M(t). \tag{3}$$

Further, the fractional intact ($P$) and damaged ($D$) protein content $[g(gDW)^{-1}]$ are described by two coupled ODEs. Intact proteins get damaged at a rate $f(t)$ and damaged proteins are repaired at a rate $r(t)$, such that

$$\frac{\mathrm{d}P(t)}{\mathrm{d}t} = -f(t)P(t) + r(t)D(t) \tag{4}$$

$$\frac{\mathrm{d}D(t)}{\mathrm{d}t} = +f(t)P(t) - r(t)D(t). \tag{5}$$

We assume the total protein fraction to be constant $P(t) + D(t) = const$, however the composition of intact and damaged proteins changes over time.
For constant rate parameters $g(t) = g$, $f(t) = f$ and $r(t) = r$ the solutions to Eq (3)-(4) can easily be obtained by calculating eigenvalues and eigenvectors.

$$M(t) = M(0) \cdot e^{gt} \tag{6}$$

$$P(t) = \frac{1}{f+r} \cdot \left[r(P(0) + D(0)) - (D(0)r - P(0)f)e^{-(f+r)t}\right] \tag{7}$$

$$D(t) = \frac{1}{f+r} \cdot \left[f(P(0) + D(0)) + (D(0)r - P(0)f)e^{-(f+r)t}\right]. \tag{8}$$

We incorporate cell division as a discrete instantaneous event in the model. Let $s \in [0.5, 1]$ denote the size (= mass) proportion of the mother cell at cell division. Then, as soon as enough biomass has been produced, $M(t_d) = s^{-1}M(0)$, the cell can divide into a mother cell and a daughter cell of sizes

$$\begin{array}{ll} \textbf{mother} & \textbf{daughter} \\ M \leftarrow sM(t_d) = M(0) & M \leftarrow (1-s)M(t_d) = (1-s)s^{-1}M(0). \end{array} \tag{9}$$

At the same time, the total fractional protein content in both compartments remains constant. Without damage retention mechanisms, we assume that also $P$ and $D$ individually remain constant across the compartments. Increasing the retention factor $re \in [0, 1]$ accounts for the asymmetric distribution of damage at cell division [29, 30], resulting in a higher fraction of damaged proteins in the mother cell compartment and a lower fraction of damaged proteins in the daughter cell compartment. To ensure that the masses in both compartments are conserved, the fraction of intact proteins is at the same time decreased or increased respectively in mother and daughter. Consequently, if at cell division the content is $P(t_d)$ and $D(t_d)$, the variables are updated according to

$$
\begin{array}{ll}
\textbf{mother} & \textbf{daughter} \\
P \leftarrow (1 - re)P(t_d) & P \leftarrow (1 + re)P(t_d) \\
D \leftarrow (1 + re)D(t_d) & D \leftarrow (1 - re)D(t_d).
\end{array}
\tag{10}
$$

### Multi-scale model suggests the trade-off between protein and ATP demand as a driver of metabolic changes during yeast replicative ageing
### Supplementary text 2: Computational guidelines

Barbara Schnitzer [*1,2], Linnea Österberg [*3], Iro Skopa [1,2], Marija Cvijovic [1,2]

March 7, 2022

[*] Authors contributed equally
[1] Department of Mathematical Sciences, Chalmers University of Technology, Gothenburg, Sweden
[2] Department of Mathematical Sciences, University of Gothenburg, Gothenburg, Sweden
[3] Department of Biology and Biological Engineering, Chalmers University of Technology, Gothenburg, Sweden

#### Contents

**1 Parameters** **2**

**2 Pseudo-code for integrated simulation** **3**

### 1 Parameters

In the lifespan simulations several parameters were fixed, according to literature and chosen with the help simulation results.

| module | description | parameter | value | unit | estimation/reference |
|---|---|---|---|---|---|
| ecFBA | inital protein content | $P_{tot}$ | 0.46 | $g(gDW)^{-1}$ | [1] |
| | fraction of included enzymes | $f$ | 0.1799 | | estimated with help of [2] |
| | enzyme saturation | $\sigma$ | 0.4592 | | adapted such that ecFBA replicates chemostat experiment [3] |
| | growth associated maintenance | $GAM$ | * | $mmol(gDW)^{-1}$ | [4, 5] |
| | maximal growth rate | | 0.35 | $h^{-1}$ | |
| | maximal acetate production | | 1.25 | $mmol(gDW \cdot h)^{-1}$ | [6] |
| | maximal glycerol production | | 1.2 | $mmol(gDW \cdot h)^{-1}$ | [6] |
| | maximal pyruvate production | | 0.1 | $mmol(gDW \cdot h)^{-1}$ | [6] |
| | maximal damage production via peroxynitrite | | 0.1 | $mmol(gDW \cdot h)^{-1}$ | |
| | maximal non-growth associated maintenance | $NGAM_{max}$ | 0.7 | $mmol(gDW \cdot h)^{-1}$ | [7] |
| | maximal superoxide production in cytosol | | $10^{-3}$ | $mmol(gDW \cdot h)^{-1}$ | needed to allow flux through damage reactions in cytosol |
| ODE | size proportion | $s$ | 0.64 | | [8] |
| | retention factor | $re$ | 0.3 | | [8, 9] |
| integrated model | threshold for glucose availability | $glc_c^{in}$ | 3.2914 | $mmol(gDW \cdot h)^{-1}$ | [5] |
| | threshold for damage production | $d_c$ | $10^{-3}$ | $mmol(gDW \cdot h)^{-1}$ | estimated with help of simulations |
| | threshold for Trx1/2 | $trx_c$ | $2 \cdot 10^{-9}$ | $mmol(gDW)^{-1}$ | estimated with help of simulations, equals two times the precision of the LP solver |
| | time step | $\delta t$ | 0.1 | $h$ | |
| | time steps between signalling and regulation | $n_{delay}$ | 5 | | |
| | flexibility in growth | $\gamma$ | 0.5 | | estimated to allow enough flexibility in enzyme reallocation during regulation |

**Table 1:** Fixed parameters and conventions used in the paper.

* The $GAM$ value is a coefficient in the stoichiometric matrix $S$ and a function of the growth rate. It increases linearly from 18 to 30 $mmol(gDW)^{-1}$ for growth rates between 0 and 0.285 $h^{-1}$ (respiration in chemostat setting), and thereafter decreases from 30 to 25 $mmol(gDW)^{-1}$ for growth rates between 0.285 and 0.4 $h^{-1}$ (fermentation in chemostat setting).

#### 2 Pseudo-code for integrated simulation

**Algorithm 1** Sketch of integrated lifespan simulation

\# INITIALISATION ##########################################

*boolean* ← initialise Boolean model from model files ”species.txt” and ”rules.txt”
*targets* ← get targets of transcription factors in the Boolean model from ”TFtargets.txt”
$\epsilon$ ← set regulation factor

*ecFBA* ← initialise ecFBA model from model file ”reducedEcYeast_XXX.mat” as a linear program
if necessary, manually curate *ecFBA* (e.g. updating bounds, additional constraints, $\sigma$, $f$)
define objective function of *ecFBA* to maximal growth

$P(0)$, $D(0)$, $M(0)$ ← set initial conditions of the ODE variables
$s$ ← set size proportion
$re$ ← set retention factor
$f_0$, $r_0$ ← set non-metabolic damage formation and repair rate
$\delta t$ ← set time step

\# SIMULATION ############################################

$t \leftarrow 0$
**while** true **do**

  \# UPDATE ecFBA PARAMETERS ##################################

  **if** deletion experiment **then**
    $e_{max,k} \leftarrow 0.0 \;\; \forall$ deleted enzymes $k$
  **end if**

  set preconditions for time-step by
  $P_{tot} \leftarrow P(t)$
  $e_{pool,max} \leftarrow f \cdot \sigma \cdot P_{tot}$
  $NGAM_{min} \leftarrow NGAM_{max} \frac{D(t)}{P(t)+D(t)}$

  \# SOLVE FOR THE FIRST TIME ################################

  solve parsimonious *ecFBA*
  **if** *ecFBA* == INFEASIBLE **then**
    break
  **end if**

  restrict growth rate with some flexibility by
  $g \leftarrow ecFBA.growthRate$
  $ecFBA.growthRate_{min} \leftarrow g \cdot (1 - \gamma)$

  $GAM \leftarrow GAM(g)$

**Algorithm 1** Sketch of integrated lifespan simulation (continued)

### REGULATE ACCORDING TO BOOLEAN SIGNALLING ##################

run *boolean* to get the logical steady state

find all transcription factors $tf$ with $boolean.tf.active == true$
match with the ones included in *targets* to get those target enzymes that are regulated

calculate ranks using
$(rank)_i \leftarrow 0$ for all enzymes $i$ in $ecFBA$
**for all** enzymes $i$ **do**
    $(rank)_i \ += 1$ for each active $tf$ that upregulates $e_i$
    $(rank)_i \ -= 1$ for each active $tf$ that downregulates $e_i$
**end for**

and update bounds on enzymes usages accordingly
**for all** enzymes i with $(rank)_i \neq 0$ **do**
    $\Delta_i \leftarrow$ calculate range of enzyme usage $e_i$ in $ecFBA$ that does not change the
        objective value (enzyme variability analysis)
    **if** $(rank)_i > 0$ **then**
        $e_{min,i} \ += \Delta_i \cdot \epsilon$
    **else if** $(rank)_i < 0$ **then**
        $e_{max,i} \ -= \Delta_i \cdot \epsilon$
    **end if**
**end for**

### SOLVE THE REGULATED MODEL ##############################

solve regulated parsimonious $ecFBA$
**if** $ecFBA ==$ INFEASIBLE **then**
    break
**end if**

**if** overexpression experiment **then**
    $e_{min/max,k} \leftarrow 1.5 \cdot e_k \ \ \forall$ overexpressed enzymes $k$
    solve updated regulated parsimonious $ecFBA$ once more
    **if** $ecFBA ==$ INFEASIBLE **then**
        break
    **end if**
**end if**

### SOLVE ODE MODEL WITH ecFBA PARAMETERS ####################

$f_m \leftarrow \sum ecFBA.damageProduction$
$g \leftarrow ecFBA.growthRate$

solve system of ODEs for $\delta t$ starting with current $M(t)$, $P(t)$ and $D(t)$
$\frac{\mathrm{d}M(t)}{\mathrm{d}t} = gM(t)$
$\frac{\mathrm{d}P(t)}{\mathrm{d}t} = -(f_m + f_0)P(t) + r_0 D(t)$
$\frac{\mathrm{d}D(t)}{\mathrm{d}t} = +(f_m + f_0)P(t) - r_0 D(t)$

**Algorithm 1** Sketch of integrated lifespan simulation (continued)

98:<br>
99: # CELL DIVISION ##########################################<br>
100:<br>
101: cell division<br>
102: **if** $M(t) \geq s^{-1}M(0)$ **then**<br>
103: $M(t) \leftarrow M(0)$<br>
104: $P(t) \leftarrow (1 - re)P(t)$<br>
105: $D(t) \leftarrow (1 + re)D(t)$<br>
106: **end if**<br>
107:<br>
108: # SAVE STEP (IF WANTED) ####################################<br>
109:<br>
110: save variables of interest (e.g. $M(t)$, $P(t)$, $D(t)$, current number of divisions, $g$, ...)<br>
111:<br>
112: # UPDATE ecFBA PARAMETERS ##################################<br>
113:<br>
114: trigger signalling with the solution fluxes and enzyme usages for next time step<br>
115: $boolean.glucose \leftarrow ecFBA.glucoseFlux > glc_c^{in}$<br>
116: $boolean.H2O2 \leftarrow ecFBA.damageProduction > d_c$<br>
117: $boolean.trx \leftarrow ecFBA.trxUsage > trx_c$<br>
118:<br>
119: unconstrain $ecFBA.growthRate$, $\mathbf{e}_{min}$ and $\mathbf{e}_{max}$ again for next time step<br>
120:<br>
121: $t \leftarrow t + \delta t$<br>
122:<br>
123: **end while**<br>
124:

One can add a time delay of the regulation by running line 50 with the Boolean model that was generated $n_{delay}$ time steps ago instead of the one from the current time step. The updates in the Boolean model from line 114-117 are then only relevant in $n_{delay}$ time steps.
